# Seed Coat Pattern QTL and Development in Cowpea (*Vigna unguiculata* [L.] Walp.)

**DOI:** 10.1101/514455

**Authors:** Ira A. Herniter, Ryan Lo, María Muñoz-Amatriaín, Sassoum Lo, Yi-Ning Guo, Bao-Lam Huynh, Mitchell Lucas, Zhenyu Jia, Philip A. Roberts, Stefano Lonardi, Timothy J. Close

## Abstract

The appearance of the seed is an important aspect of consumer preference for cowpea (*Vigna unguiculata* [L.] Walp.). Seed coat pattern in cowpea has been a subject of study for over a century. This study makes use of newly available resources, including mapping populations, a reference genome and additional genome assemblies, and a high-density single nucleotide polymorphism genotyping platform, to map various seed coat pattern traits to three loci, concurrent with the *Color Factor* (*C*), *Watson* (*W*), and *Holstein* (*H*) factors identified previously. Several gene models encoding proteins involved in regulating the later stages of the flavonoid biosynthesis pathway have been identified as candidate genes, including a basic helix-loop-helix gene (*Vigun07g110700*) for the *C* locus, a WD-repeat gene (*Vigun09g139900*) for the *W* locus and an E3 ubiquitin ligase gene (*Vigun10g163900*) for the *H* locus. A model of seed coat development, consisting of six distinct stages, is described to explain some of the observed pattern phenotypes.

## 1 Introduction

Cowpea (*Vigna unguiculata* [L.] Walp.) is a diploid (2n = 22) warm season legume which is primarily grown and serves as a major source of protein and calories in sub-Saharan Africa. Further production occurs in the Mediterranean Basin, southeast Asia, Latin America, and the United States. Just over 7.4 million metric tonnes of dry cowpeas were reported worldwide in 2017 (FAOSTAT, 2019), though these numbers do not include Brazil, Ghana, and some other relatively large producers. Most of the production in sub-Saharan Africa is by smallholder farmers in marginal conditions, often as an intercrop with maize, sorghum, or millet (Ehlers and Hall, 1997). Due to its high adaptability to both heat and drought and its association with nitrogen fixing bacteria, cowpea is a versatile crop (Ehlers and Hall, 1997; Boukar et al., 2018).

The most common form of consumption is as dry grain. The seeds are used whole or ground into flour (Singh, 2014; Tijjani et al., 2015). Seed coat pattern is an important consumer-related trait in cowpea. Consumers make decisions about the quality and presumed taste of a product based on appearance (Jaeger et al., 2018; Kostyla et al.,1978). Cowpea displays a variety of patterns, including varied eye shapes and sizes, Holstein, Watson, and Full Coat pigmentation, among others (Figure 1). Each cowpea production region has preferred varieties, valuing certain color and pattern traits above others for determining quality and use. In West Africa consumers pay a premium for seeds exhibiting certain characteristics specific to the locality, such as lack of color for use as flour or solid brown for use as whole beans (Herniter et al., 2019; Langyintuo et al., 2003; Mishili et al., 2009). In the United States consumers prefer varieties with tight black eyes, commonly referred to as “black-eyed peas” (Fery, 1985).

**Figure 1.**
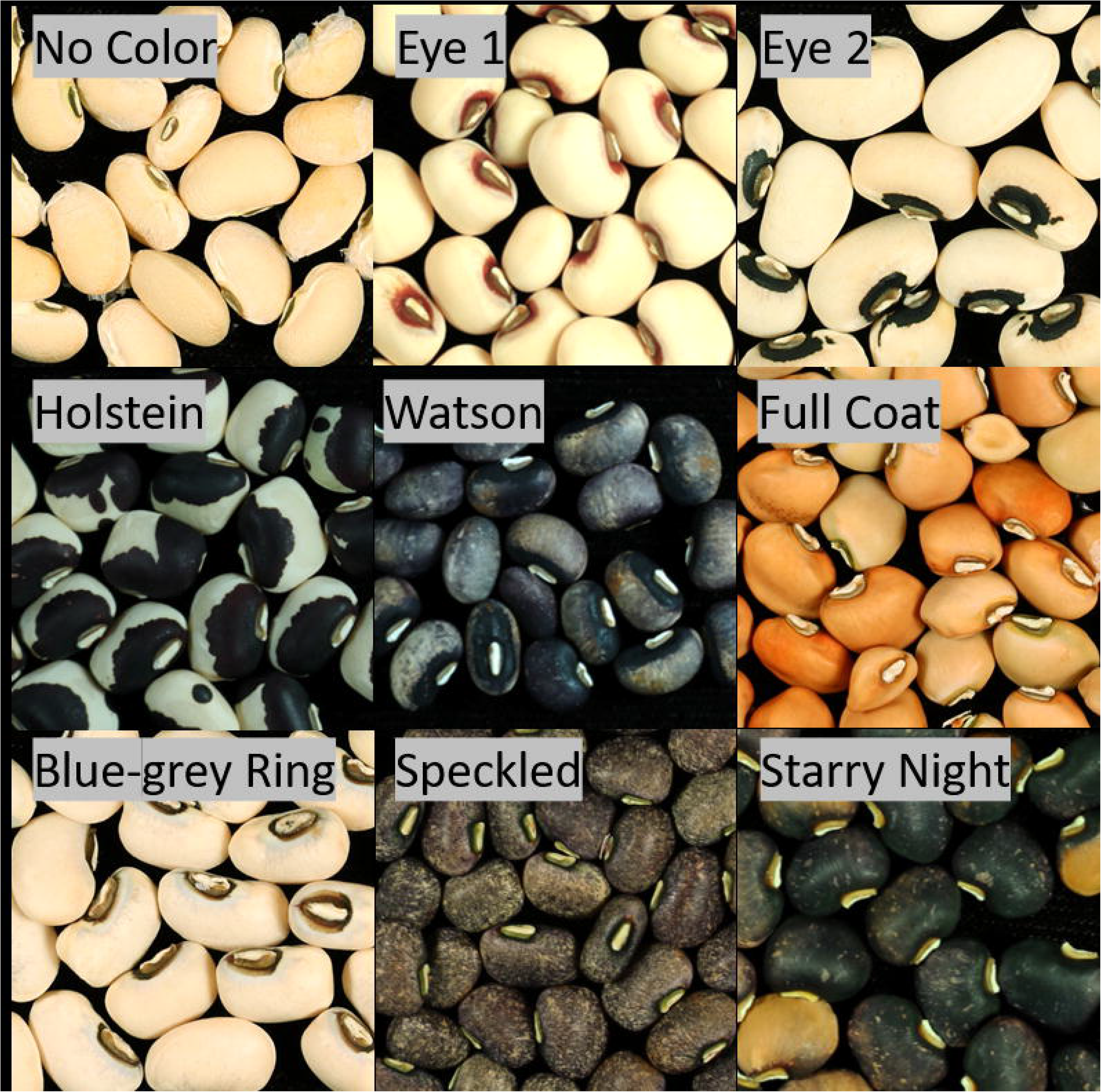
Seed coat pattern traits. Images of lines from various populations demonstrating the phenotypes which were scored as part of this study.

Seed coat traits in cowpea have been studied since the early 20th century, when Spillman (1911) and Harland (1919), reviewed by Fery (1980), explored the inheritance of factors controlling seed coat color and pattern. In a series of F2 populations Spillman (1911) and Harland (1919) identified genetic factors responsible for color expression, including “*Color Factor*” (*C*), “*Watson*” (*W*), “*Holstein-1*” (*H-1*), and “*Holstein-2*” (*H-2*). A three-locus system controlling seed coat pattern was established by Spillman and Sando (1930) and was confirmed by Saunders (1960) and Drabo et al. (1988), though “*O*” was used in place of “*C*.”

A genotyping array for 51,128 single nucleotide polymorphisms (SNP) was recently developed for cowpea (Muñoz-Amatriaín et al., 2017) which offers opportunities to improve the precision of genetic mapping. Numerous biparental populations have been used to map major quantitative trait loci (QTL) for various traits, including root-knot nematode resistance (Santos et al., 2016), domestication-related traits (Lo et al., 2018), and black seed coat color (Herniter et al., 2018) and to develop consensus genetic maps of cowpea (Lucas et al., 2011; Muchero et al., 2009; Muñoz-Amatriaín et al., 2017). In addition, new populations have been developed for higher-resolution mapping including an eight-parent Multi-parent Advanced Generation Inter-Cross (MAGIC) population containing 305 lines (Huynh et al., 2018). A reference genome sequence of cowpea (Lonardi et al., 2019; phytozome.net) and genome assemblies of six additional diverse accessions (Muñoz-Amatriaín et al., 2019) have been produced recently. Here, we make use of these resources to map a variety of seed coat pattern traits, determine candidate genes, and develop a model for genetic control of seed coat pattern. Additionally, we posit a developmental pattern for the cowpea seed coat to explain some of the observed variation.

## 2 Materials and Methods

### 2.1 Plant Materials

Ten populations were used for mapping: an eight-parent MAGIC population containing 305 lines (Huynh et al., 2018), four biparental recombinant inbred line (RIL) populations, and five F2 populations. Descriptions of each pattern discussed below can be found in Section 2.3 and examples can be seen in Figure 1.

One biparental population consisted of 87 RILs developed at the University of California, Riverside (UCR), derived from a cross between California Blackeye 27 (CB27), which has a black Eye 2 pattern, and IT82E-18, also known as “Big Buff” (BB), which has a brown Full Coat pattern (Muchero et al., 2009). The second biparental RIL population consisted of 80 RILs developed at UCR derived from a cross between CB27 and IT97K-556-6 (556), which has a brown Full Coat pattern (Huynh et al., 2015). The third biparental RIL population consisted of 101 RILs developed at UCR, derived from a cross between California Blackeye 46 (CB46), which has a black Eye 2 pattern, and IT93K-503-1 (503), which has a brown Eye 1 pattern (Pottorff et al., 2014). The fourth biparental RIL population consisted of 76 RILs developed at UCR and at the International Institute for Tropical Agriculture in Nigeria, derived from a cross between 524B, which has a black Eye 2 pattern, and IT84S-2049 (2049), which has a brown Eye 1 pattern (Menéndez et al., 1997). The F2 populations were developed at UCR as part of this work. Two F2 populations, consisting of 176 and 132 individuals, were developed from independent crosses between CB27 and Bambey 21 (B21), which has the No Color phenotype. One F2 population, consisting of 143 individuals, was developed from a cross between B21 and California Blackeye 50 (CB50), which has a black Eye 2 pattern. Two F2 populations, consisting of 175 and 119 individuals, were developed from independent crosses between Tvu-15426, which has a purple Full Coat pattern, and MAGIC014, a line developed as part of the MAGIC population but not included in the final population, which has a black Watson pattern.

To temporally describe seed coat development four accessions were examined: CB27, MAGIC059, Sanzi, and Sasaque. CB27 is described above. MAGIC059 has the Starry Night pattern in black and purple and is one of the lines included in the MAGIC population. Sanzi has a Speckled pattern in black and purple. Sasaque has the Full Coat pattern in red and purple.

### 2.2 SNP genotyping and data curation

DNA was extracted from young leaf tissue using the Qiagen DNeasy Plant Mini Kit (Qiagen, Germany). A total of 51,128 SNPs were assayed in each sample using the Illumina Cowpea iSelect Consortium Array (Illumina Inc., California, USA; Muñoz-Amatriaín et al., 2017). Genotyping was performed at the University of Southern California Molecular Genomics Core facility (Los Angeles, California, USA). The same custom cluster file as in Muñoz-Amatriaín et al. (2017) was used for SNP calling. In the F2 populations the extracted DNA was bulked by phenotype, with DNA from 20 individuals combined in each genotyped sample.

For the MAGIC population, SNP data and a genetic map were available from Huynh et al. (2018). The map included 32,130 SNPs in 1,568 genetic bins (Huynh et al., 2018). For the biparental RIL populations, SNP data and genetic maps for the CB27 by BB and the CB46 by 503 populations were available from Muñoz-Amatriaín et al. (2017), and SNP data and a genetic map were available for the 524B by 2049 population from Santos et al. (2018). The CB27 by 556 genetic map was created using MSTMap (Wu et al., 2008). The CB27 by BB genetic map included 16,566 polymorphic SNPs in 977 genetic bins (Muñoz-Amatriaín et al., 2017); the CB27 by 556 genetic map contained 16,284 SNPs in 2604 bins; the CB46 by 503 genetic map contained 16,578 SNPs in 683 bins (Muñoz-Amatriaín et al., 2017); the 524B by 2049 genetic map contained 14,202 SNPs in 933 bins (Santos et al., 2018). For each F2 population, SNPs were filtered to remove non-polymorphic loci between the respective parents. The number of markers used for each population is as follows: the two CB27 by B21 populations, 8,550 SNPs (Supplementary Table 1); the B21 by CB50 population, 8,628 SNPs (Supplementary Table 2); the two Tvu-15426 by MAGIC014 populations, 20,010 SNPs (Supplementary Table 3).

### 2.3 Seed coat phenotyping

Phenotype data for seed coat traits were collected by visual examination of the seeds. The scored phenotypic classes consisted of No Color, Eye 1, Eye 2, Holstein, Watson, and Full Coat (Figure 1). No Color indicates no pigmentation present on the seed coat. Eye 1 consists of a loose eye in the shape of a teardrop with spots of color outside the eye on the wider side. Eye 2 consists of a tight eye in the shape of two wings with no pigment observed outside the edge of the eye. Holstein consists of an eye with a defined edge and additional spots of pigmentation spread over the seed coat up to almost completely covering the coat. Watson consists of an eye with an indefinite edge. Full Coat consists of pigment completely covering the seed coat. Two of the lines used for observing seed coat development had other seed coat patterns than those mapped. MAGIC014 had the Starry Night pattern, which consists of incomplete pigmentation covering the entire seed. Sanzi had the Speckled pattern, which consists of small dots of pigment covering the seed coat. Seeds with a paler brown color are often difficult to distinguish between the Eye 1 and Watson patterns. The MAGIC population was scored for Eye 1, Eye 2, Holstein, Watson, and Full Coat patterns (Supplementary Table 4). The CB27 by BB (Supplementary Table 5) and CB27 by 556 (Supplementary Table 6) biparental RIL populations were scored for Eye 2, Holstein, Watson, and Full Coat patterns. The CB46 by 503 (Supplementary Table 7) and 524B by 2049 (Supplementary Table 8) biparental RIL populations were scored for Eye 1, Eye 2, Holstein, Watson, and Full Coat patterns. The CB27 by B21 and B21 by CB50 F2 populations were scored for the No Color and Eye 2 patterns. The Tvu-15426 by MAGIC014 F2 populations were scored for the Watson and Full coat patterns.

For mapping purposes, each observed pattern was scored individually and mapped independently with scores assigned as “1” indicating presence of the trait and a “0” indicating absence. For example, a line expressing the Eye 1 pattern would be scored as “1” for the Eye 1 trait and “0” for all other traits. Pattern phenotypes are mutually exclusive. As the Eye 1 pattern appears to be epistatic towards the *H* and *W* loci, any lines with the Eye 1 phenotype were scored as missing data for other seed coat phenotypes to avoid biasing the mapping. This was the case in all populations other than the MAGIC population, as the mpMap script could not operate with such an extent of missing data. In the MAGIC population, for traits other than Eye 1 (Eye 2, Holstein, Watson, and Full Coat), individuals with the Eye 1 phenotype were scored as “0” instead of as missing data since marking too many lines as missing data caused r/mpMap to fail.

### 2.4 Segregation Ratios

Expected segregation ratios reported in Table 2 were determined based on the type of population, parental and F1 phenotypes. For example, the F2 populations were expected to segregate in a 3:1 ratio for traits controlled by single genes with complete dominant/recessive relationships, while the biparental RIL populations were expected to segregate in a 1:1 ratio. Expected segregation ratios were tested by chi-square analysis.

For the MAGIC population, based on how the population was constructed (Huynh et al., 2018) it was assumed that each fully homozygous parent had a roughly 1/8 probability to pass its genotype at a particular locus to a given RIL. For example, at the *C* locus, three parents (IT84S-2049, IT89KD-288, and IT93K-503-1) express the Eye 1 phenotype and are proposed to have a *C*_*1*_*C*_*1*_ genotype, while the other five parents are proposed to have a *C*_*2*_*C*_*2*_ genotype. Based on this, a given line in the population is expected to have a 3/8 probability of having a *C*_*1*_*C*_*1*_ genotype and a 5/8 probability of have a *C*_*2*_*C*_*2*_ genotype. At the *W* and *H* loci, one parent (CB27) is proposed to have the *H*_*0*_*H*_*0*_ and *W*_*0*_*W*_*0*_ genotypes, while the other seven parents are proposed to have the *W*_*1*_*W*_*1*_ and *H*_*1*_*H*_*1*_ genotypes. Based on this, a line should have a 1/8 probability of having the *W*_*0*_*W*_*0*_ and a 1/8 probability of having the *H*_*0*_*H*_*0*_ genotype. By multiplying the probabilities at each locus, the probability of a given genotype can be determined using the following equation:

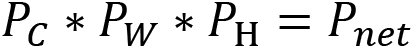

Where P_C_ is the probability of a given allele at the *C* locus, P_W_ is the probability of a given allele at the *W* locus, P_H_ is the probability of a given allele at the *H* locus, and P_net_ is the probability of a given genotype. For example, the probability of a *C*_*2*_*C*_*2*_*H*_*1*_*H*_*1*_*W*_*0*_*W*_*0*_ genotype, which would have a Holstein phenotype would be 35/512 ([5/8]*[7/8]*[1/8]). The above method results in a predicted 192:5:35:35:245 phenotypic ratio for the Eye 1 (*C*_*1*_*C*_*1*_), Eye 2 (*C*_*2*_*C*_*2*_*H*_*0*_*H*_*0*_*W*_*0*_*W*_*0*_), Holstein (*C*_*2*_*C*_*2*_*H*_*1*_*H*_*1*_*W*_*0*_*W*_*0*_), Watson (*C*_*2*_*C*_*2*_*H*_*0*_*H*_*0*_*W*_*1*_*W*_*1*_), and Full Coat (*C*_*2*_*C*_*2*_*H*_*1*_*H*_*1*_*W*_*1*_*W*_*1*_) patterns, respectively.

### 2.5 Trait mapping

Trait mapping was achieved with different methods for each type of population. In the MAGIC population, the R package “mpMap” (Huang and George, 2011) was used as described by Huynh et al. (2018). The significance cutoff values were determined through 1000 permutations, resulting in a threshold of *p* = 8.10E-05 [−log10(*p*) = 4.09]. Due to the high number of markers in the genotype data, imputed markers spaced at 1 cM intervals were used.

In the biparental RIL populations, the R packages “qtl” (Broman et al., 2003) and “snow” (Tierney et al., 2015) were used as in Herniter et al. (2018). Briefly, probability values were assigned to each SNP using a Haley-Knott regression, tested for significance with 1000 permutations, and marker effects were determined using a hidden Markov model.

For the F2 populations, the genotype calls of each bulked DNA pool in the population were filtered to leave only the markers known to be polymorphic between the parents, and these were then sorted based on physical positions in the pseudochromosomes available from Phytozome (Lonardi et al. 2019; phytozome.net). Each population’s genotype was then examined visually in Microsoft Excel for areas where the recessive bulk was homozygous, and the dominant bulk was heterozygous. Duplicated populations were examined in conjunction.

### 2.6 Determining haplotype blocks

Once significant regions were established through mapping analysis, the overlapping area shared between the four biparental RIL populations was examined to determine the minimal area where all four biparental populations had overlapping haplotype blocks. SNPs located in the hotspots of pseudochromosomes Vu07, Vu09, and Vu10 were examined visually in Microsoft Excel for regions of identity within phenotypic groups. SNPs located in the hotspots which had been removed during trait mapping due to high levels of missing data were added back as presence/absence variations and segregated similar to nucleotide polymorphisms.

### 2.7 Determining candidate genes

Genes were examined within each minimal haplotype block. Gene expression data (Yao et al., 2016), from the cowpea reference genome (IT97K-499-35), which has a black Eye 1 (*C*_*1*_*C*_*1*_) pattern available from the Legume Information System (legumeinfo.org) were examined for expression in developing seed tissue. Genes encoding proteins known to be involved in regulation of the flavonoid biosynthesis pathway were prioritized.

### 2.8 Determining allelic series

Dominance relationships were determined by examining the phenotypes of several F1 progeny in addition to segregation ratios in the F2 populations. Crosses were made between CB27 and three lines from the CB27 by BB population (BB-090, BB-113, and BB-074). Seeds from these F1 plants were visually examined for seed coat patterns. CB27/BB-090 seeds had a Watson pattern (*C*_*2*_*C*_*2*_*H*_*0*_*H*_*0*_*W*_*1*_*W*_*1*_), CB27/BB-113 seeds had a Holstein pattern (*C*_*2*_*C*_*2*_*H*_*1*_*H*_*1*_*W*_*1*_*W*_*1*_), and CB27/BB-074 seeds had a Full Coat pattern (*C*_*2*_*C*_*2*_*H*_*1*_*H*_*1*_*W*_*1*_*W*_*1*_). An additional cross was available from the early development of the MAGIC population, where the phenotype of the seed coat on seeds from a maternal *C*_*1*_*C*_*2*_ heterozygote was Full Coat. IT84S-2246 (Full Coat, *C*_*2*_*C*_*2*_*H*_*1*_*H*_*1*_*W*_*1*_*W*_*1*_) was crossed with IT93K-503-1 (Eye 1, *C*_*1*_*C*_*1*_*H*_*1*_*H*_*1*_*W*_*1*_*W*_*1*_) to yield this *C*_*1*_*C*_*1*_*H*_*1*_*H*_*1*_*W*_*1*_*W*_*1*_ maternal parent.

### 2.9 Comparing sequence variation

The genome sequences of the candidate genes from each of five genome sequences (the reference genome sequence and four additional genome assemblies) and about 3 kb of upstream sequence were compared using A plasmid Editor (ApE; jorgensen.biology.utah.edu/wayned/ape/). Transcription factor binding sites were predicted in the upstream regulatory region of each gene using the binding site prediction function available from the Plant Transcription Factor Database (Jin et al., 2017; planttfdb.cbi.pku.edu.cn/). The species input was *Vigna radiata* (mung bean), as a map of cowpea was unavailable. The cowpea reference sequence is of IT97K-499-35. Among the additional sequenced genomes, CB5-2 has the Eye 2 pattern (*C*_*2*_*C*_*2*_), Suvita-2 has the Full Coat pattern (*C*_*2*_*C*_*2*_*H*_*1*_*H*_*1*_*W*_*1*_*W*_*1*_), Sanzi has a Speckled pattern, and UCR779 has the Full Coat pattern (*C*_*2*_*C*_*2*_*H*_*1*_*H*_*1*_*W*_*1*_*W*_*1*_). See Section 2.3 for pattern descriptions and Figure 1 for examples.

A larger set of SNPs (about 1 million), discovered from whole-genome shotgun sequencing of 37 diverse accessions (Muñoz-Amatriaín et al., 2017; Lonardi et al. 2019)), was available from Phytozome (phytozome.net). Among the 37 accessions, 28 had phenotype data available. These lines were examined for variations in the SNP selection panel that were in the gene-coding and regulatory regions of the candidate genes.

### 2.10 Correlation test

The 28 lines from the SNP selection panel with phenotype and genotype data available were tested for correlation in R, using the native “cor.test” function. For input, the phenotype was recorded as “+1” for accessions with the Eye 1 (*C*_*1*_*C*_*1*_) phenotype and “-1” for those without. The genotype was recorded as “+1” for accessions matching the reference genotype, “−1” for the alternate homozygote, and “0” for the heterozygote (Supplementary Table 9).

### 2.11 Seed color development

The four accessions for which pattern development was recorded (CB27, MAGIC059, Sanzi, and Sasaque) were grown in a greenhouse at the University of California, Riverside (Riverside, California; 33.97° N 117.32° W) at a constant temperature of about 32°C from March through May 2018. Three plants were used for each accession. Upon flowering, each flower was tagged with the date it opened. The flowers were permitted to self-fertilize. For each day after the flower opened, beginning on the second day, on each of the three test plants a pod was collected until no more green pods were observed.

Seeds from each collected pod were photographed using a Canon EOS Rebel T6i at a 90° angle under consistent lighting conditions. The length of the most advanced seed within the pod was measured using ImageJ (imagej.nih.gov). A developmental scale from 0 to 5 was designed based on the visual observations of the spread of pigmentation (see Results). Each photograph was scored using this scale.

## 3. Results

### 3.1 Phenotypic data and segregation ratios

Phenotypic data and proposed genotypes for each parent in the observed populations can be found in Table 1. A summary of the phenotypic data, along with predicted segregation ratios, chi-square values, and probability can be found in Table 2.

**Table 1.** Parental phenotypes and expected genotypes of the examined populations.

| Population | Population type | Parent | Phenotype | Proposed Genotype |
| --- | --- | --- | --- | --- |
| MAGIC | 8-Parent RIL | California Blackeye 27 | Eye 2 | $C_2C_2W_0W_0H_0H_0$ |
| | | IT00K-1263 | Full Coat | $C_2C_2W_1W_1H_1H_1$ |
| | | IT82E-18 | Full Coat | $C_2C_2W_1W_1H_1H_1$ |
| | | IT84S-2049 | Eye 1 | $C_1C_1W_1W_1H_1H_1$ |
| | | IT84S-2246 | Full Coat | $C_2C_2W_1W_1H_1H_1$ |
| | | IT89KD-288 | Eye 1 | $C_1C_1W_1W_1H_1H_1$ |
| | | IT93K-503-1 | Eye 1 | $C_1C_1W_1W_1H_1H_1$ |
| | | SuVita 2 | Full Coat | $C_2C_2W_1W_1H_1H_1$ |
| CB27 by BB | Biparental RIL | California Blackeye 27 | Eye 2 | $C_2C_2W_0W_0H_0H_0$ |
| | | IT82E-18 | Full Coat | $C_2C_2W_1W_1H_1H_1$ |
| CB27 by 556 | Biparental RIL | California Blackeye 27 | Eye 2 | $C_2C_2W_0W_0H_0H_0$ |
| | | IT97K-556-6 | Full Coat | $C_2C_2W_1W_1H_1H_1$ |
| CB46 by 503 | Biparental RIL | California Blackeye 46 | Eye 2 | $C_2C_2W_0W_0H_0H_0$ |
| | | IT93K-503-1 | Eye 1 | $C_1C_1W_1W_1H_1H_1$ |
| 524B by 2049 | Biparental RIL | 524B | Eye 2 | $C_2C_2W_0W_0H_0H_0$ |
| | | IT84S-2049 | Eye 1 | $C_1C_1W_1W_1H_1H_1$ |
| CB27 by B21 | F2 | California Blackeye 27 | Eye 2 | $C_2C_2W_0W_0H_0H_0$ |
| | | Bambey 21 | No Color | $C_0C_0W_0W_0H_0H_0$ |
| B21 by CB50 | F2 | Bambey 21 | No Color | $C_0C_0W_0W_0H_0H_0$ |
| | | California Blackeye 50 | Eye 2 | $C_2C_2W_0W_0H_0H_0$ |
| Tvu-15426 by MAGIC014 | F2 | Tvu-15426 | Full Coat | $C_2C_2W_1W_1H_1H_1$ |
| | | MAGIC014 | Watson | $C_2C_2W_1W_1H_0H_0$ |

**Table 2.** Phenotypes, segregation ratios, and probability values for the tested populations.

| Population<br>(# of lines) | Eye 1 | Eye 2 | Holstein | Watson | Full<br>Coat | No<br>Color | Pred. Seg. Ratio | X<sup>2</sup> | Probability |
| --- | --- | --- | --- | --- | --- | --- | --- | --- | --- |
| MAGIC<br>(305) | 121 | 0 | 21 | 13 | 141 | -- | 192:5:35:35:245 | 6.41 | 0.17 |
| CB27 by<br>Big Buff<br>(87) | -- | 20 | 28 | 16 | 23 | -- | 1:1:1:1 | 3.53 | 0.32 |
| CB27 by<br>556 (80) | -- | 14 | 30 | 17 | 19 | -- | 1:1:1:1 | 7.30 | 0.06 |
| CB46 by<br>503 (101) | 49 | 12 | 17 | 8 | 15 | -- | 4:1:1:1:1 | 3.73 | 0.44 |
| 524B by<br>2049 (76) | 47 | 5 | 8 | 6 | 10 | -- | 4:1:1:1:1 | 5.82 | 0.21 |
| CB27 by<br>B21 A<br>(176) | -- | 129 | -- | -- | -- | 47 | 3:1 | 0.27 | 0.60 |
| CB27 by<br>B21 B<br>(132) | -- | 88 | -- | -- | -- | 44 | 3:1 | 4.89 | 0.027 |
| B21 by<br>CB50 (143) | -- | 112 | -- | -- | -- | 31 | 3:1 | 0.84 | 0.36 |
| Tvu-15426<br>by<br>MAGIC014<br>A (175) | -- | -- | -- | 44 | 131 | -- | 1:3 | 0.0019 | 0.97 |
| Tvu-15426<br>by<br>MAGIC014<br>B (120) | -- | -- | -- | 27 | 93 | -- | 1:3 | 0.40 | 0.53 |

### 3.2 Identification of loci controlling seed coat pattern

A total of 35 SNP loci were identified using different methods for each population type (see Materials and Methods for details) and were concentrated on three chromosomes: Vu07 (*C* locus), Vu09 (*H* locus), and Vu10 (*W* locus). Mapping results can be found in Supplementary Table 10. The overlapping mapping results allowed a narrowing of the area examined for candidate genes.

### 3.3 Determination of minimal haplotype blocks

Following trait mapping, all called SNPs on chromosomes Vu07, Vu09, and Vu10 were examined for minimal haplotype blocks in the overlapping significant regions in the four biparental RIL populations. On Vu07 (*C* locus) the minimal haplotype block was between 2_12939 and 2_09638 (228,331 bp) and contained ten genes. On Vu09 the minimal haplotype block was between 2_33224 and 2_12692 (166,724 bp) and contained seventeen genes. On Vu10 the minimal haplotype block was between 2_12467 and 2_15325 (120,513 bp) and contained eleven genes. The list of candidate genes can be found in Supplementary Table 11 and on Phytozome (Lonardi et al. 2019; phytozome.org) The minimal haplotype block regions can be found in Supplementary Table 12.

### 3.4 Identification of candidate genes

A predominant candidate gene was identified at each locus based on high relative expression in the developing seeds (Supplementary Figure 1) and a review of the literature on the regulation of the flavonoid biosynthesis pathway (see Discussion for details). This led to the determination of a single major candidate gene on each of Vu07, Vu09, and Vu10. Each of the candidate genes belongs to a class which is known to be involved in transcriptional control of the later stages of flavonoid biosynthesis. No Color, Eye 1, and Full Coat mapped to an overlapping area on Vu07, where the gene *Vigun07g110700*, encoding a basic helix-loop-helix protein, was noted as a strong candidate gene. Eye 2, Holstein, Watson, and Full Coat mapped to a similar area on Vu09, where the gene *Vigun09g139900*, encoding a WD-repeat gene, was noted as a strong candidate gene. Eye 1, Eye 2, Holstein, Watson, and Full Coat mapped to an overlapping area on Vu10, where the gene *Vigun10g163900*, encoding an E3 ubiquitin ligase protein with a zinc finger, was noted as a strong candidate gene.

### 3.5 Determination of allelic series

Segregation ratios indicated the dominance of *H*_*1*_ over *H*_*0*_ (*Holstein* locus, Figure 2E, Gii), *W*_*1*_ over *W*_*0*_ (*Watson* locus, Figure 2Gi), *C*_*2*_ over *C*_*0*_ (*Color Factor* locus, Figure 2F), and *C*_*2*_ over *C*_*1*_ (*Color Factor* locus, Figure 2Giv). The dominance relationship between the *C*_*1*_ and C_0_ alleles could not be determined from these data.

**Figure 2.**
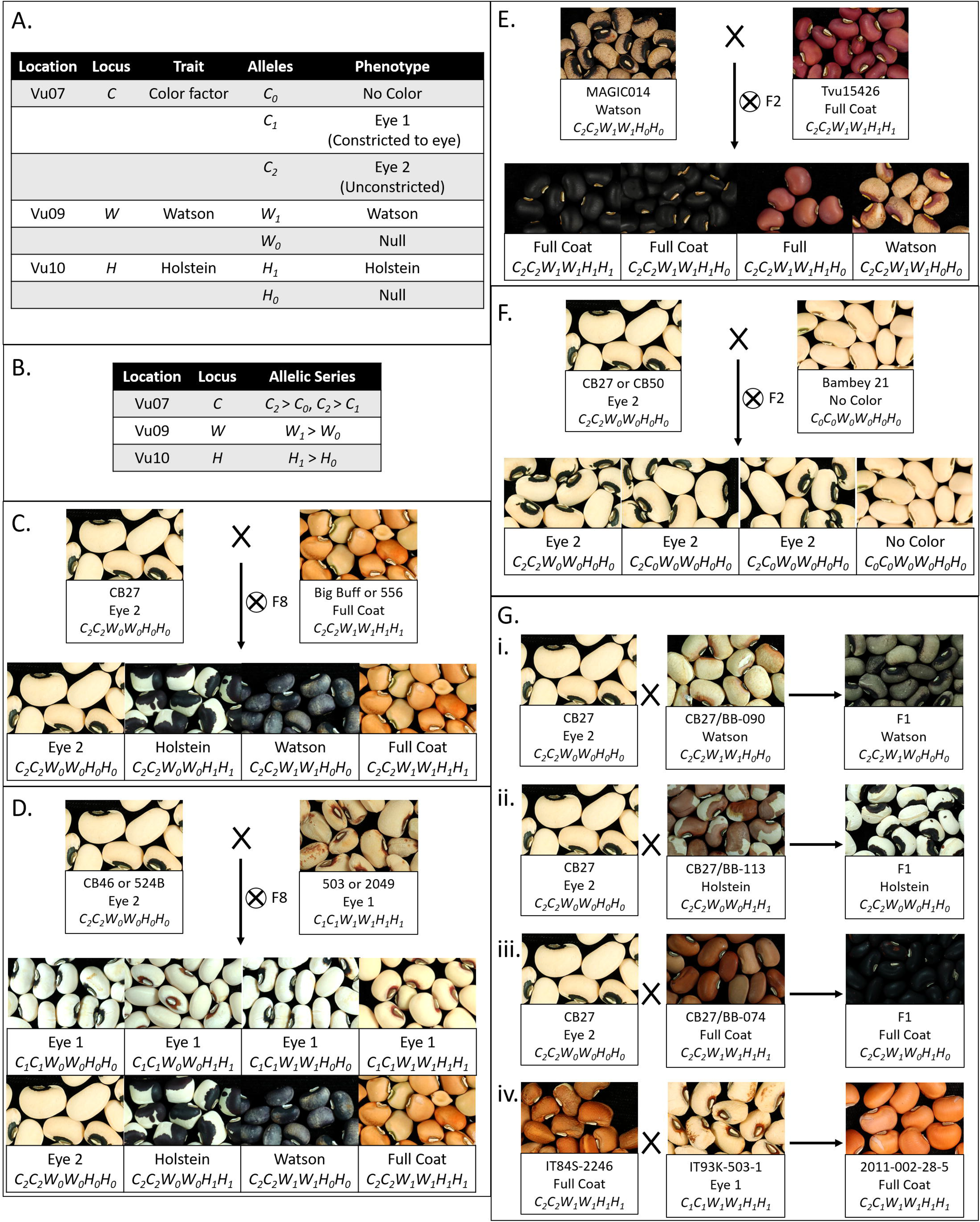
Interaction of seed coat pattern loci. (A) Table displaying the pattern loci identified in mapping, their locations, the trait encoded, alleles identified, and phenotypes. (B) Table displaying the allelic series and relative dominance of alleles. (C) Segregation patterns for the CB27 by BB and CB27 by 556 F8 populations. (D) Segregation patterns for the CB46 by 503 and 524B by 2049 F8 populations. (E) Segregation pattern for the Tvu-15426 by MAGIC014 F2 populations. (F) Segregation pattern for the CB27 by B21 and B21 by CB50 F2 populations. (G) Phenotype of seeds from the F1 plants resulting from a series of crosses (i) Cross between CB27 and line from the CB27 by BB population with a Watson pattern, resulting in Watson pattern. (ii) Cross between CB27 and a line from the CB27 by BB population a Holstein pattern, resulting in Holstein pattern. (iii) Cross between CB27 and a line from the CB27 by BB population with a Full Coat pattern, resulting in a Full Coat pattern. (iv) Cross between IT84S-2246 and IT93K-503-1 from the early development of the MAGIC population, resulting in a Full Coat pattern in the seed coats on seeds of the F1 maternal parent.

### 3.6 Sequence comparisons of candidate genes

Multiple sequence alignments for each of the three candidate genes and regulatory regions (~3 kb upstream of the transcription start site) revealed SNPs and small insertions or deletions (Supplementary Datasets 1, 2, and 3). None of the variants in the transcript sequence were predicted to cause changes in the amino acid sequence.

The regulatory region of *Vigun07g110700* (*C* locus candidate gene) showed a C/T SNP variation between the reference genome and the four other genome sequences on Vu07 at 20,544,306 bp. The reference genome has a T at this position while the other four sequences have a C. Transcription factor binding site prediction from the Plant Transcription Factor Database (planttfdb.cbi.pku.edu.cn/) indicated that this variation constitutes either a WRKY binding site in the C allele or an ERF binding site in the T allele. Of the 28 accessions in the SNP selection panel, eleven expressed the Eye 1 (*C*_*1*_) pattern and 17 did not. Twenty accessions had a CC genotype, six had a TT genotype, and two had a TC genotype. The correlation test gave an estimated correlation value of 0.75, with a *p*-value of 3.51E-06, indicating significant correlation between the genotype and phenotype values such that this SNP is a reliable marker for distinguishing between the *C*_*1*_ (Eye 1) and the *C*_*2*_ (Eye 2) alleles. Two of the 28 lines had the No Color (*C*_*0*_) phenotype, but had the CC genotype, indicating that this SNP is not a good marker for the *C*_*0*_ allele (for a possible explanation see Discussion). The regulatory region of *Vigun09g139900* (*W* locus candidate gene) showed a C/T variation between the reference genome and CB5-2 against the other three genome sequences on Vu09 at 30,207,722 bp. This SNP was not included in the list from the SNP selection panel and so could not be examined like the previous SNP. Transcription factor binding site prediction did not indicate that the site was a target for any transcription factor in either form. The upstream regulatory region of *Vigun10g163900* (*H* locus candidate gene) did not have any distinguishing variation.

### 3.7 Stages of color development

A model of seed coat development has been formulated consisting of six stages based on the spread of pigmentation. In Stage 0, there is no color on the seed coat. In Stage 1, color appears at the base of the hilum. In Stage 2, color appears around the hilum. In Stage 3, color begins to spread along the outside edges of the seed. In Stage 4, color begins to fill in on the edges of the testa. In Stage 5, the color has completely developed to the mature level. After Stage 5 the pod and seeds begin to desiccate. Of the observed varieties, only Sasaque and Sanzi completed all six stages. MAGIC059 reached Stage 4, while CB27 only reached Stage 2. No seeds in Stage 0 were observed for Sasaque. Images of each tested variety at various stages can be seen in Figure 3. Color development was associated with seed size; the pigmentation spread as the seeds grew larger.

**Figure 3.**
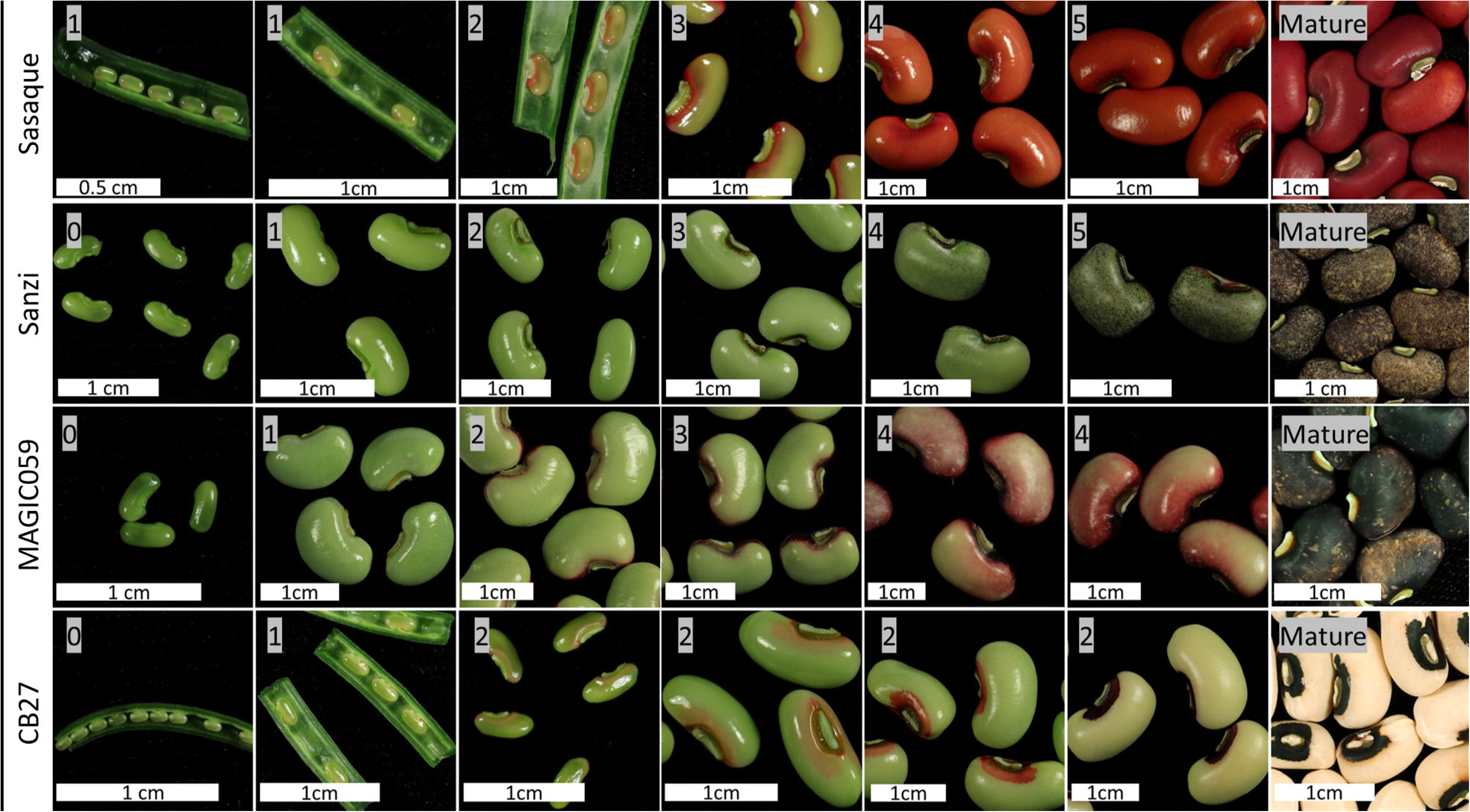
Seed coat color development. Images showing the development the seed and the spread of pigmentation.

## 4 Discussion

### 4.1 Segregation ratios and epistatic interaction of seed coat pattern loci

Segregation ratios and dominance data (Table 2, Figure 2) in the tested populations were consistent with a three gene system with simple dominance and epistatic interactions that matches the *C* (*Color Factor*), *W* (*Watson*), and one of the *H* (*Holstein*) factors identified by Spillman (1911) and Harland (1919). In brief, the *C* locus encodes a “constriction” factor while the *W* and *H* loci encode distinct “expansion” factors. The *C* locus is the primary locus controlling seed coat pattern. Pigmentation may be not visible (No Color, *C*_*0*_), constrained to an eye (Eye 1, *C*_*1*_), or distributed throughout the seed coat (Eye 2, Holstein, Watson, or Full Coat, *C*_*2*_). The extent of distribution is modified by the *H* and *W* loci, whose contribution is visible only with an unconstrained allele (*C*_*2*_) at the *C* locus. In the presence of *Holstein* (*H*_*1*_) and absence of *Watson* (*W*_*0*_), a Holstein pattern is expressed. Conversely, in the presence of *Watson* (*W*_*1*_) and absence of *Holstein* (*H*_*0*_), a Watson pattern is expressed. In combination, the *Watson* (*W*_*1*_) and *Holstein* (*H*_*1*_) factors result in the Full Coat phenotype.

Based on the above proposed allelic series, an individual with the *C*_*0*_*C*_*0*_ genotype will express the No Color pattern, regardless of the genotypes at the *W* and *H* loci, and an individual with the *C*_*1*_*C*_*1*_ genotype will express the Eye 1 pattern, regardless of the genotypes at the *W* and *H* loci. However, when not constricted by a *C*_*0*_ or *C*_*1*_ allele (having the *C*_*2*_ allele) the “expansion” factors can be observed. An individual with the *C*_*2*_--*W*_*0*_*W*_*0*_*H*_*1*_--genotype expresses the Holstein pattern, while and individual with the *C*_*2*_--*W*_*1*_--*H*_*0*_*H*_*0*_ genotype expresses the Watson pattern. An individual with the *C*_*2*_--*W*_*1*_--*H*_*1*_--genotype, with both “expansion” factors, expresses the Full Coat pattern. An individual with the *C*_*2*_--*W*_*0*_*W*_*0*_*H*_*0*_*H*_*0*_ genotype expresses the Eye 2 pattern. In this latter case the eye pattern is observed despite the unconstricted *C*_*2*_ allele due to the absence of the “expansion” factors. Based on this model, the CB27 by BB and CB27 by 556 populations segregate at the *W* and *H* loci (Figure 2C), while the MAGIC, CB46 by 503, and 524B by 2049 populations segregate at all three loci (Figure 2D). Similarly, the Tvu-15426 by MAGIC014 populations segregate at the *W* locus (Figure 2E) and the CB27 by B21 and B21 by CB50 populations segregate at the *C* locus (Figure 2F).

An additional pattern phenotype of Blue-grey Ring was noted in some of the tested populations. Blue-grey Ring consists of a pale ring of bluish-grey surrounding the eye (Figure 1). It appears only with the Eye 1 (*C*_*1*_) phenotype but is not always present when the phenotype is Eye 1 (*C*_*1*_). The Blue-grey Ring phenotype may represent another (fourth) allele at the *C* locus, or it may result from a combination of the *C*_*1*_ (Eye 1) allele and other pigmentation genes. However, from other unpublished work on seed coat color there does not appear to be a strict correlation between seed coat color and presence of the Blue-grey Ring. Further research is required to clarify the basis of the Blue-grey Ring phenotype.

### 4.2 Pattern traits QTL overlap

Several regions of the genome are hotspots for seed coat pattern traits (Supplementary Table 11). These correspond to locations of genetic factors identified by Spillman (1911) and Harland (1919), who identified four factors controlling seed coat patterning: *Color Factor* (*C*), *Watson* (*W*), *Holstein-1* (*H-1*), and *Holstein-2* (*H-2*). The present data suggest the presence of only one *Holstein* locus or that the two loci are very closely linked in the tested populations. To avoid possible confusion, the *Holstein* locus discussed here is simply termed “*H*.”

The major QTL and regions of interest for No Color and Eye 1 are clustered in an overlapping region on Vu07, suggesting that the “constriction” factor at locus *C* is at that position with allelism at the locus. Mapping results from the Tvu-15426 by MAGIC014 F2 populations indicate that the *H* locus is on Vu10. Additional evidence for the *H* locus being located on Vu10 comes from Wu et al. (2019), who identified the *Anasazi* locus (equivalent to the cowpea *H* locus) on chromosome 10 of common bean, which is homologous to Vu10 (Lonardi et al., 2019). While none of the biparental F2 populations segregated solely for the *W* locus, the identification of the *C* locus on Vu07 and the *H* locus on Vu10 must, by process of elimination, identify the location of the *W* “expansion” locus on Vu09.

### 4.3 Seed coat pattern is due to failure to complete the normal color developmental program

It was noted that the varieties with the Full Coat pattern at maturity followed the developmental pattern described in Section 3.7 and shown in Figure 3 to completion. In contrast, varieties which do not display the Full Coat pattern appear to have color development arrested at certain points. This is most obvious in CB27 (Eye 2, *C*_*2*_), where color development proceeds only to Stage 2. It is likely that other varieties which have distinct eye sizes proceed to varied stages of development. For example, varieties with the No Color (*C*_*0*_) phenotype would not proceed past Stage 0. However, the three gene model presented here does not explain every seed coat pattern. An example is the pattern observed in mature Sanzi seed, which exhibits a Speckled black and purple seed coat (see Section 2.3 for a description and Figure 1). According to this analysis, Sanzi completes all six stages of seed coat development, indicating that the Speckled pattern is controlled separately. A biparental RIL population, consisting of lines derived from a cross between Sanzi and Vita 7, which has a brown Full Coat pattern (*C*_*2*_*C*_*2*_*W*_*1*_*W*_*1*_*H*_*1*_*H*_*1*_), was used for mapping the black seed coat color; there was a perfect correlation between black seed coat color and the Speckled pattern (Herniter et al., 2018). This indicates that genetic control of the Speckled pattern is colocalized with black seed coat color and may be an allele at the *Bl* locus, which is located on Vu05.

Further research is needed to determine if all cowpea accessions follow the pattern observed in the four tested lines shown in Figure 3. It may be that each of the observed stages of seed coat pigmentation development is controlled by a different gene, and that failures of normal gene function cause the observed variation in patterning. Evidence for this model is furnished by the noted developmental pattern of the seed coats where development appears to be arrested at Stage 2 in CB27, which expresses the Eye 2 (*C*_*2*_) pattern, and at Stage 4 in MAGIC059, which expresses the Starry Night pattern (see Section 2.3 for a description and Figure 1). The mechanism by which this occurs is not elucidated here and requires further research. Transcriptome data could be gathered for the seed coat at each developmental stage. The currently available transcriptome data (Yao et al., 2016; legumeinfo.org) used whole seeds at specific days post flowering and do not distinguish between transcripts in the seed coat and those in the embryo or cotyledons, and further do not separate transcripts by developmental stage.

### 4.4 Candidate gene function

The later steps in flavonoid biosynthesis are controlled by a transcription factor complex composed of an R2-R3 MYB protein, a basic helix-loop-helix protein (bHLH), and a WD-repeat protein (WD40; Xu et al., 2015). E3 Ubiquitin ligases (E3UL) are believed to negatively regulate this complex (Shin et al., 2015). The color and location (leaf, pod, seed coat) of the pigmentation are determined by expression patterns (Wu et al., 2003, Iorizzo, 2018). Candidate genes on Vu07 (*C* locus) and Vu09 (*W* locus) encode a bHLH and WD40 protein, respectively. A candidate gene on Vu10 (*H* locus) encodes an E3UL protein. This information lends itself to a model in which *Vigun07g110700* (bHLH) serves as a “master switch” controlling the extent of pigmentation constriction while *Vigun09g139900* (WD40) and *Vigun10g163900* (E3UL) act as “modulating switches” controlling the type of expanded pattern, altering the effect of the pathway to result in the observed Holstein and Watson patterns (Figure 4). The R2-R3 MYB directs the DNA binding of the complex, with expression of different genes in different tissues resulting in the observed color and location of the pigments. For example, MYB genes identified by Herniter et al. (2018) are required for black seed coat and purple pod tip color. Further, *Vigun07g110700* (bHLH) was identified as a candidate gene controlling flower color in cowpea by Lo et al. (2018), indicating a possible dual function of the gene. Indeed, Harland (1919) noted that a lack of pigment in the flower was often associated with a lack of pigment in the seed coat. Finally, homologs of *Vigun07g110700* have been identified in other legumes as Mendel’s *A* gene controlling flower color in *Pisum sativum* (Hellens et al., 2010) and as the *P* gene in *Phaseolus vulgaris* (McClean et al., 2018).

**Figure 4.**
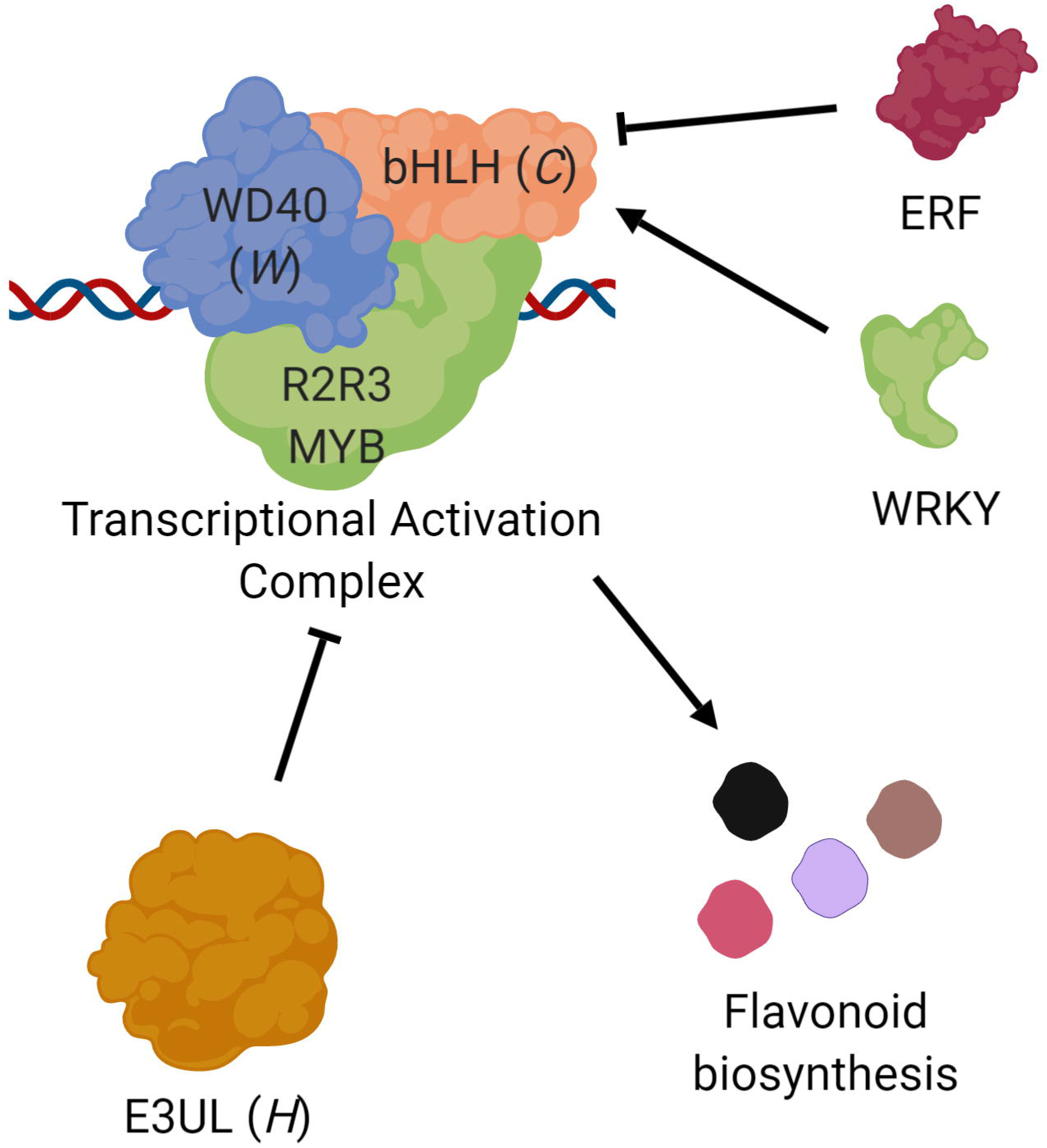
Proposed roles of the *C*, *W*, and *H* genes. Transcription of flavonoid biosynthesis pathway genes are controlled by a complex composed of three types of proteins (Xu et al., 2015), a basic helix-loop-helix protein (bHLH; e.g., *Vigun07g110700*, *C* locus), a WD-repeat protein (WD40; e.g., *Vigun09g139900*, *W* locus), and an R2R3 MYB transcription factor. This complex is in turn negatively regulated by an E3 Ubiquitin ligase (E3UL; e.g., *Vigun10g163900*, *H* locus). Sequence comparisons suggest that bHLH transcription may be controlled by ERF and WRKY proteins. The observed seed coat pattern phenotypes are a result of different alleles and expression patterns.

Two R2R3 *MYB* genes (*Vigun10g165300* and *Vigun10g165400*) are located only 110 kb downstream of *Vigun10g163900* (*H* locus candidate gene). However, these fall outside of the haplotype blocks identified in the CB27 by BB and CB27 by 556 populations, indicating that they are not the source of the observed phenotypic variation. However, there may be interaction between one or both of these MYBs and the E3UL responsible for the Holstein pattern; this hypothesis could be investigated through additional research.

The observed C/T SNP variation in the regulatory sequence of *Vigun07g110700* (bHLH) at 20,544,306 bp constitutes a difference between a WRKY binding site in the *C*_*2*_ (Eye 2) allele versus an ERF binding site in the *C*_*1*_ (Eye 1) allele. WRKY proteins are positive regulators of seed coat pigment biosynthesis in Arabidopsis (Lloyd et al., 2017) while ERF proteins negatively regulate the same pathway (Matsui et al., 2008). This SNP could be used as a genetic marker to distinguish between the *C*_*1*_ and *C*_*2*_ alleles. The lack of correlation between an observed marker and the *C0* (No Color) allele may be caused by other variants, such as a small deletion interrupting gene function, which has been shown in *Phaseolus vulgaris* (McClean et al., 2018). Such a variation would not be detected by the genotyping platform used for this study. Similarly, the observed C/T SNP variation in the regulatory region of *Vigun09g139900* at 30,207,722 bp could be used as a marker to distinguish between the *W*_*0*_ (not Watson) and *W*_*1*_ (Watson) alleles, despite not necessarily being the cause of the observed phenotypic variation. No single variation was identified for *Vigun10g163900* alleles. However, haplotype blocks determined from the biparental RIL populations can be used for future breeding efforts. Two SNPs which fall within the genome sequence of *Vigun10g163900* segregate with the phenotype in the biparental RIL populations. At 2_24359, the lines with the *H*_*0*_ (not Holstein) allele have an A genotype and the lines with the *H*_*1*_ (Holstein) allele have a G genotype. At 2_24360, the lines with the *H*_*0*_ (not Holstein) allele have an A and the lines with the *H*_*1*_ (Holstein) allele have a C. Future research is needed to develop more perfect markers for the three loci.

## 5 Abbreviations

2049: IT84S-2049
503: IT93K-503-1
556: IT97K-556-6
B21: Bambey 21
BB: Big Buff (IT82E-18)
bHLH: basic helix-loop-helix
*C*: *Color Factor*
CB27: California Blackeye 27
CB46: California Blackeye 46
CB50: California Blackeye 50
E3UL: E3 Ubiquitin ligase
GWA: Genome-Wide Association
*H*: *Holstein*
MAGIC: Multiparent Advanced Generation InterCross
QTL: quantitative trait locus
RIL: recombinant inbred line
SNP: single nucleotide polymorphism
UCR: University of California Riverside
*W*: *Watson*
WD40: WD-repeat

## 6 Acknowledgements

This manuscript has been released as a Pre-Print in bioRxiv (Herniter et al., 2019). The authors thank Amy Litt for helpful discussion and guidance on pattern development; Eric Castillo and Sabrina Phengsy for assistance with seed photography; Steve Wanamaker for assistance in the analysis of the various genome sequences. This study was supported by the Feed the Future Innovation Lab for Climate Resilient Cowpea (USAID Cooperative Agreement AID-OAA-A-13-00070), the National Science Foundation BREAD project “Advancing the Cowpea Genome for Food Security” (NSF IOS-1543963) and Hatch Project CA-R-BPS-5306-H.

## 7 Author Contributions Statement

I.H. performed all trait mapping, statistical analysis, and interpretation. R.L. performed analysis of the seed coat development. M.M. assisted in trait mapping and provided SNP data. Sa.L. assisted in trait mapping. Y.G. extracted DNA for genotyping. B.H. provided the MAGIC population and its genotypic information. M.L. performed crosses used for allelic series analysis. Z.J. assisted in statistical analysis. P.R. and T.C. provided guidance and access to population and genetic resources. St.L. assisted with the SNP selection panel data. T.C. assisted I.H. with the writing.

*The authors declare that the research was conducted in the absence of any commercial or financial relationships that could be construed as a potential conflict of interest*.

## 8 Contribution to the Field

Seed coat pattern is an important consumer-related trait. Consumers make decisions about the quality, value, and use of products based on visual traits. As such, it is important for breeders to understand the genetic bases of these traits to facilitate efforts to produce improved varieties that meet market preferences. Previous research, dating back to the early twentieth century, first reported genetic factors controlling cowpea seed coat pattern. With access to new resources, including genome sequences, mapping populations, and advanced genetic markers, here we clarify the inheritance of and interactions between major loci controlling seed coat patterns. Specifically, this includes three candidate genes for control of seed coat pattern and possible genetic markers that can be used for breeding purposes. In addition, we propose a model of seed coat development to explain much of the observed variation. Our findings advance the understanding of the genetic control of seed coat pattern in cowpea and provide actionable results that can be applied in breeding programs.

## 9 Data Availability Statement

All datasets [SNPs] for this study are included in the manuscript and the supplementary files.

---

Supplementary Figure 1. Relative expression levels of the candidate genes. TPM, Transcripts per million; dap, days after pollination. Data retrieved from legumeinfo.org.

## Notes

#### Summary of Updates

Figure 2 has been updated with new information regarding the dominance of the C2 allele over the C1 allele. The C3 allele is nut supported by the evidence and has been removed. Authors and author affiliations have been updated.

