## Supplementary figures and images for "Seed Coat Pattern QTL and Development in Cowpea (*Vigna unguiculata* [L.] Walp.)"

### Supplementary Figure 1

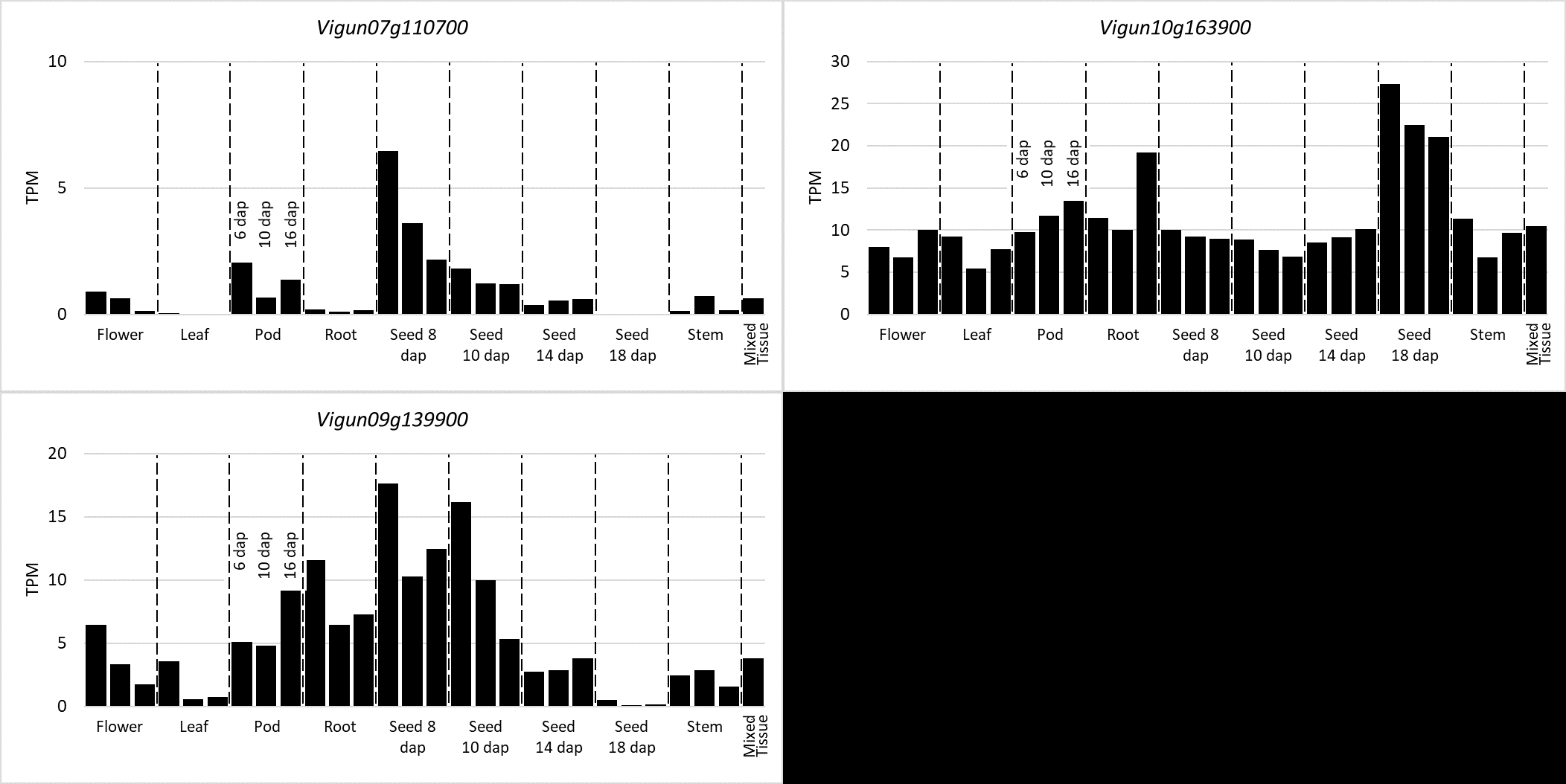
